# Immunoglobulin T genes in Neopterygii

**DOI:** 10.1101/2020.05.21.108993

**Authors:** Serafin Mirete-Bachiller, David N. Olivieri, Francisco Gambón-Deza

## Abstract

In teleost fishes there are three immunoglobulin isotypes named immunoglobulin M (IgM), D (IgD) and T (IgT). IgT has been the last to be described and is considered a teleosts-fish specific isotype. From the recent availability of genome sequences of fishes, an in-depth analysis of Actinopterygii immunoglobulin heavy chain genes was undertaken. With the aid of a bioinformatics pipeline, a machine learning software, CHfinder, was developed that identifies the coding exons of the CH domains of fish immunoglobulins. Using this pipeline, a high number of such sequences were obtained from teleosts and holostean fishes. IgT was found in teleost and holostean fishes that had not been previously described. A phylogenetic analysis reveals that IgT CH1 exons are similar to the IgM CH1. This analysis also demonstrates that the other three domains (CH2, CH3 and CH4) were not generated by recent duplication processes of IgM in Actinopterygii, indicating it is an immunoglobulin with an earlier origin.

## 1. Introduction

Some 550 million years ago the adaptive immune system arose coinciding with the appearance of the non-jawed vertebrates (Agnathans) (Cooper & Alder, 2006). The adaptive response began when antigens were recognized by specific receptors on lymphocytes. In Agnathans, such receptors are encoded by a family of genes first identified in the lamprey variable lymphocyte receptors (VLRs), while in jawed vertebrates (gnathostomes), these were the B and T lymphocyte receptors (BCR and TCR, respectively). Many features of the VLR system are analogous to BCR and TCR, although a notable difference is that the VLR system is made up of highly diverse leucine-rich repeats (LRR), whereas in gnathostomes the antigenic receptors have immunoglobulin domains (Pancer et al., 2004).

Immunoglobulin genes emerged with the gnathostomes (Flajnik & Kasahara, 2010). These genes are made up of multiple exons that code for domains of approximately 100 amino acids and with a sequence and structure that is called the immunoglobulin domain (Peterson et al., 1972; Edelman et al., 1969). These domains were already found in proteins of invertebrate species, encoding for membrane proteins with cellular communication functions and some of them participate in the innate immune response (Sun et al., 1990; Zhang et al., 2004). An organization of these genes occurred in jawed vertebrates, which gave rise simultaneously to the immunoglobulin genes (both for the heavy and light chain), the genes for T lymphocyte receptors (for both TCR *α/β* as for the TCR *γ/δ*), and the genes for the major histocompatibility complex.

Today’s cartilaginous fish have what is believed to be the oldest genes tructure of all immunoglobulins (IgLs). Genes in these animals that code for the variable (V) and constant (C) regions that appear in multiple repeated V-C tandem arrangements. Bony fishes have a differentiated structure with separate V and C gene groups. These structures evolved such that in order to have a valid mRNA coding for an immunoglobulin chain, V and C genes must be joined together. Thereafter, this process was maintained throughout all posterior vertebrate lines.

Different immunoglobulin isotypes have been described in gnathostomes. With only some exceptions, IgM and IgD are present in all jawed vertebrates, appearing approximately 500 million years ago. IgM has been stable throughout evolution, while IgD undergone great plasticity with respect to the number of CH domains and the amount of splice variants found (Ohta & Flajnik, 2006). Apart from these two Igs, other isotypes have been described in vertebrates. In amphibians and reptiles, two new isotypes were found that are not present in fishes (IgY and IgA(X)) and in mammals are been found 5 isotypes (IgM, IgD, IgG, IgE and IgA) (Gambon-Deza et al., 2009; Zhao et al., 2009).

In the 1990s, IgM was cloned and characterized in *Ictalurus punctatus* and *Gadus morhua* (Ghaffari & Lobb, 1989; Bengtén et al., 1991). Amongst all the immunoglobulins, IgM in bony fishes has the highest concentration in serum. In these species, it has some peculiarities with respect to its mammalian orthologs. First, it is tetrameric and does not possess a J chain, so polymerization occurs by interchain disulfide bonds (Castro & Flajnik, 2014). Also, by means of an alternative splicing mechanism, the membrane form of IgM consists of only three domains, however this does not prevent it from interacting with CD79 (Ross et al., 1998; Sahoo et al., 2008). IgM can be formed as monomers, dimers, and trimers; it was suggested that the degree of polymerization of IgM is related to maturation affinity (Ye et al., 2010).

IgD in Teleostei was first described in *Ictalurus punctatus* (Wilson et al., 1997). Between Teleostei species, there is a high variability in the number of constant domains. The C*μ*1 exon that contains the region coding the cysteine residue which binds to the light chains, is spliced with the first C *δ* exon. This immunoglobulin has been described in several teleost fish species (Hordvik et al., 1999; Stenvik & Jørgensen, 2000). The secreted form of IgD has been identified in *Ictalurus punctatus* and *Takifugu rubripes* (Hikima et al., 2011). In *Ictalurus punctatus*, the secreted form lacks the VH regions and can bind to basophils, mediated by an Fc receptor that induces pro-inflammatory cytokines production (Chen et al., 2009).

In 2005, two independent groups described a new immunoglobulin isotype in teleosts, with four constant domains in *Danio rerio* and *Oncorhynchus mykiss*, which they named IgZ and IgT respectively. In the same year, an isotype with two domains was described in *Takifugu rubripes* that they called IgH (Danilova et al., 2005; Hansen et al., 2005; Savan et al., 2005b). However, these immunoglobulins were shown to be orthologs to each other and whose consensus was later called IgT (Gambón-Deza et al., 2010). Later, IgT was described in more teleosts specie and found to have a variable number of constant domains, ranging from 2 to 4. Also, fishes can have more than one gene for IgT (See Table S2). However, IgT is not found in all teleosts; for example, catfish and medaka lack IgT (Bengtén et al., 2002; Magadán-Mompó et al., 2011). This isotype occurs as monomers in serum and as tetramers in mucous (Zhang et al., 2011). Studies of *Cyprinus carpio* suggested an important role of IgT in mucosal immunity (Ryo et al., 2010).

Because previous work has only found IgT in Teleostei, a widely accepted conclusion was that it was specific to species of this order. Here, we provide evidence that, in fact, IgT has its origins prior to the appearance of Teleostei. Specifically, we describe its presence in Holostei, indicating that the origin of this Ig isotype dates back to the late Permian with the appearance of the Neopterygii subclass.

## 2. Methods

### 2.1. Sequence Data

We studied the presence of immunoglobulin genes of bony fishes present from genome assemblies available from the https://vgp.github.io/ and NCBI repositories. The gene finding program CHfinder, developed by our group, was used to search and annotate immunoglobulin exons from chromosomal regions in these assemblies.

### 2.2. Bioinformatics Tools

CHfinder is based on a multiclass neural network machine learning classifier that was trained to identify exons coding for the CH domains of fish immunoglobulins. For the training process, domain sequences of IgM (CH1, CH2, CH3 and CH4), IgD (CH1, CH1B, CH2, CH3, CH4, CH5 and CH6) and IgT (CH1, CH2, CH3 and CH4), together with random background sequences constituted the training set. The resulting neural network has an accuracy of 99.99 % for identifying IG domains.

The gene identification workflow is as follows. First, a BLAST (tblastn) is used as a coarse grained filter by using a query consisting of an artificially constructed sequence, concatenating representative IgT, IgD, and IgM into a single sequence. The Tblastn query was performed against the target genome, producing a hit table - those sequences in the genome, together with their locations, having similarity to the query sequence. The coordinates of the hits are used to extract a nucleotide region, with extra padding of 1000 nucleotides at both ends. From these extracted nucleotide regions, the CHfinder identifies candidate exon reading frames, defined by the AG and GT start/stop motifs and having a min/max nucleotide length of 240nt and 450nt, respectively. Additionally, the reading frames were restricted to those that were a multiple of 3 and do not contain stop codons within the reading frame. Even with these restrictions, a large number of candidate exon sequences are obtained result that are then subjected to the machine learning prediction step of CHfinder.

Exon prediction and classification in CHfinder is done by first transforming the amino acid (AA) sequences into an integer feature vector that spatially preserves the AA ordering. Also, the transformation acts to direct training on extreme ends of each exon. In this way, each AA sequence is first converted into a sequence of 80 amino acids, by extracting and concatenating the 40 amino acids from front/back ends of the sequence coding from a hypothetical exon. These sequences are then converted to numerical “one-hot” based feature vectors. In this one-hot scheme, each amino acid in the 80aa-long exon is converted to 20 integers, all 0 except 1 for the AA in question; thus, a single exon will have a feature vector of 80 × 20 = 1600 integers. For supervised machine learning, the training set consists of known sequences with their respective IG domain label, and a set of random background sequences for null discrimination, in a ratio of 3:1 to labelled sequences.

While genome assemblies exist for many fish species, there are still several species that do not yet have published genomes. One such example is *Amia calva*. To explore the presence or absence of IgT in this species where a genome is not available, we used raw sequence records (SRA) of bowfin gills (SRX661027) from the NCBI repository. To construct transcripts, Trinity (a de novo RNA-seq transcript assembler) was used (Grabherr et al., 2011). Once assembled, the nucleotide sequences were transformed into amino acids using TransDecoder that identifies candidate coding regions within transcript sequences, such as those generated by de novo RNA-Seq transcript assembly using Trinity (Haas et al., 2013). These bioinformatics steps were performed in Galaxy (Afgan et al., 2018).

### 2.3. Phylogenetic studies

To align the IgM and IgT sequences obtained with CHfinder, the SEAVIEW (Gouy et al., 2009) and Jalview (Waterhouse et al., 2009) programs were used together with the clustal *ω* algorithm (Sievers & Higgins, 2014). Due to the heterogeneity of the genomic sources, we manually deleted the V and stem regions, in order to only retain the immunoglobulin constant domains. From this curation process, a Fasta file with all the sequences was generated (included in the Supplementary Material). Using Galaxy, sequences were aligned with MAFFT, a multiple alignment program for amino acid or nucleotide sequences (Galaxy Version 7.221.3) (Katoh & Standley, 2013). Alignments were done using the BLOSUM62 matrix. Maximum likelihood phylogenetic trees were constructed from the amino acid sequences by FastTree (Galaxy Version 2.1.10 + galaxy1) (Price et al., 2010) using the LG protein evolution model (Le & Gascuel, 2008) + CAT. Finally, the graphical representations of the trees were made using the online program iTOL (Letunic & Bork, 2019).

## 3. Results

IgT has been described in Teleostei. Indeed, this immunoglobulin is observed in several of these species, but in others it is absent. This lack of consistency throughout Teleostei raises questions about the origins and functions of IgT. Given the availability of genomes and transcriptomes from a broad sampling of species from this order, we can finally obtain a more complete catalogued the IgT prevalent in this order.

The three immunoglobulin classes found in fish are IgM, IgD, and IgT. The exons coding for the CH domains and the deduced immunoglobulin genes identified by the CHfinder program are given in Table S1. With few exceptions, all fish have genes for IgM, IgD, and most for IgT. Representative sequences for the IgT were used to search the Fish-T1K (Sun et al., 2016) repository that contains transcriptome sequences for 124 fish species, shown in Table S3. Additionally, a bibliographic search was carried out since 2005 to identify the different IgT that have been published in teleosts (see Table S2). These results indicate that the IG constant chain locus is quite unstable, having undergo duplications in some species and having large variability in the number of genes between species. For example, one species has up to 35 constant region genes (*Labrus bergylta*), while another (*Guania wildenowi*) has no genes at all.

By studying the genomes of teleost fishes, we have confirmed previous observations - most species in this order possess genes for IgT, while some do not. Nonetheless, a pattern that explain the absence of this gene remains elusive. In addition, we found genes for IgT in the Holostei clade that have not been described previously.

### 3.1. IgT in Holostei

Holostei is one of the two infraclasses of the Neopterygii subclass (Teleostei, is the other), and it comprises two orders, the Amiiformes with a single representative and which is considered a living fossil *Amia calva* and the Lepisosteiformes with two genera Atractosteus and Lepisosteus, with 7 extant species between both. The *Lepisosteus oculatus* (spotted gar) genome is available from the NCBI with a chromosomal assembly level (accession number GCA 000242695.1). A search for the immunoglobulin constant regions was performed using CHfinder. Three scaffolds containing Ig exons coding for CH domains were identified: NC 023183.1 (LG5 chromosome) contains four exons coding for CHs domains of IgM and 12 exons for the IgD domains, NW 006270078.1 has a gene with four exons for the CH domains (first identified as CH1M (similar to CH1 of IgM), while the rest of the domains were identified as CH2T, CH3T and CH4T). The scaffold NW 006270215.1 has another gene with 3 exons coding for domains: the first exon, is identified as the CH1 of IgM (CH1M), while the others were identified as CH3 of IgT (CH3T) and CH4 of IgT (CH4T) (See Figure 1).

**Figure 1:**
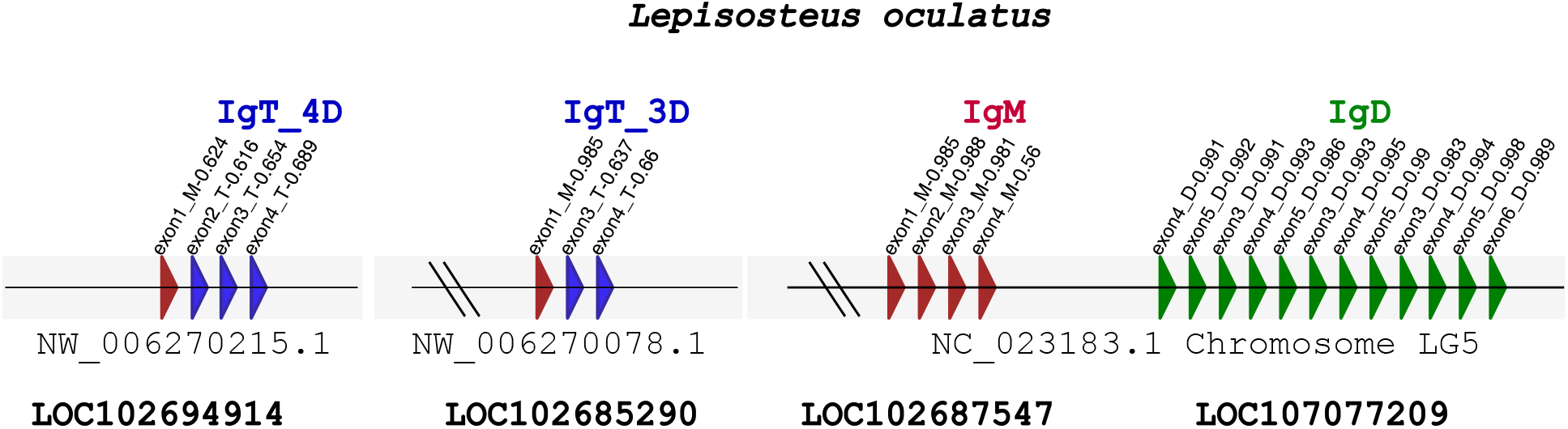
A schematic representation of the immunoglobulin genes found in the *Lepisosteus oculatus* genome. NC 023183.1 corresponds to chromosome LG5 containing the genes for IgM (red) and IgD (green). NW 006270215.1 and NW 006270078.1 are not yet assigned to chromosome and contain the exons for two different IgTs, one for 4 domains and the other for 3 domains, respectively.

Candidate IgT sequences in the spotted gar genome (IgT4D, an IgT consisting of four domains, and IgT3D, an IgT consisting of three domains) were used to search for orthologs in other Holostei species. Because the genome of *Amia calva* (bowfin) is not available, we searched for the presence of these exons in the RNA-seq transcript (SRX661027). Sequences of four CH domain orthologous to the spotted gar IgT4D were found. In the Fish-T1K repository, we identified another IgT4D (*Atractosteus spatula*) in the scaffold WHYR15010142 A C1141006. Additionally, these genes were searched in the genomes and transcripts of Polypteriformes and Acipenseriformes (firsts Actinopterygii) without finding IgT orthologues. In the amino acid sequences of these IgT in Holostei found cysteine residues are important in the formation of intra-domain disulfide bridges. Additionally, the CH1 domain contains the necessary Cys residue to form the disulfide bridge between IgH chain and IgL chain (see Fig 2).

**Figure 2:**
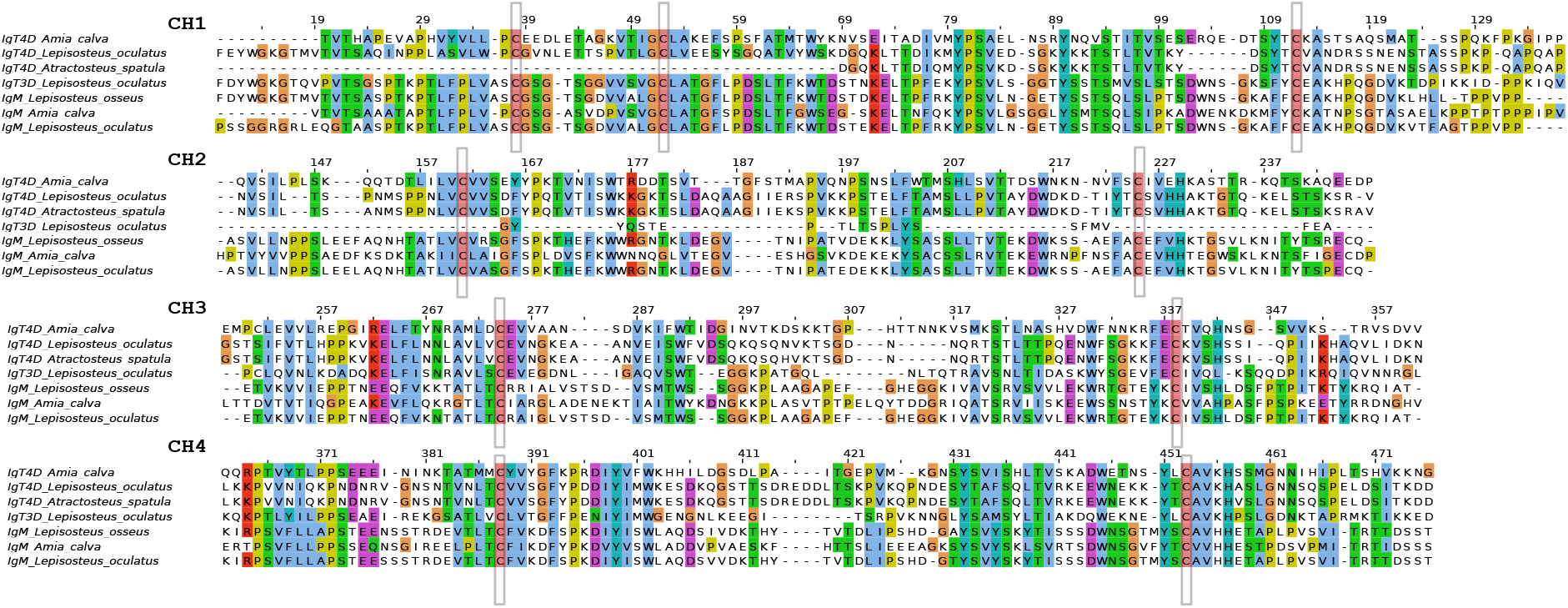
Alignment of the IgT and IgM immunoglobulin domains sequences of Holosteos. The graph was obtained with Jalview. The IgT sequences from *Lepisosteus oculatus* were deduced from the sequenced genome using CHfinder, while those of *Amia calva* and *Atractosteus spatula* were obtained from RNA-seq transcriptomes. The IgM sequences of *Lepisosteus oculatus* was obtained from CHfinder, while the IgM of *Amia calva* and *Lepisosteus osseus* were obtained from GenBANK with accession number AAC59687 and AAC59688, respectively.

### 3.2. IgT in Teleostei

CHfinder allowed us to identify 136 candidate IgT genes from the 73 Actinopterygii (see Table S1). In the Fish-T1K repository, 57 IgT sequences from 57 fish species were identified. Additionally, a bibliographical search identified 24 articles that made reference to IgT sequences in 24 teleost fishes. The obtained data set was processed manually in order to eliminate those sequences that were incomplete or repeated. Finally, we obtained 142 complete IgT sequences and 88 IgM sequences from 88 Actinopterygii species. With these sequences, a comparative phylogenetic study was undertaken.

Evolutionary relationships between the IgT and the other immunoglobulins IgM, IgW, and IgNAR can be observed from comparative phylogenetic trees of these sequences. Additionally, the sequences from Elasmobranchii were also included since their origin dates prior to that of Actinopterygii. Thus, using the appearance of the Elasmobranchii as a reference, three clades can be be observed in the resulting tree: one for Neopterygii IgT, one for Actinopterygii IgM, and a third clade encompassing the Elasmobranchii immunoglobulins. Within the IgM clade, the appearance of two Neopterygii IgT subclades is observed, one is for the 2-domain IgT found in Cichliform fish, and the other is for the IgT3D found in *Lepisosteus oculatus*. These two anomalies may be explained by the high homology of its CH1 with CH1M (both domains have quasi-identical sequences), together with the decrease in the number of domains present.

The clade corresponding to the majority of IgT sequences is unique and does not share a common node with IgM from Actinopterygii. This suggests an early origin, thereby discounting the notion of an IgM duplication in the evolution towards Actinopterygii (see Fig 3).

**Figure 3:**
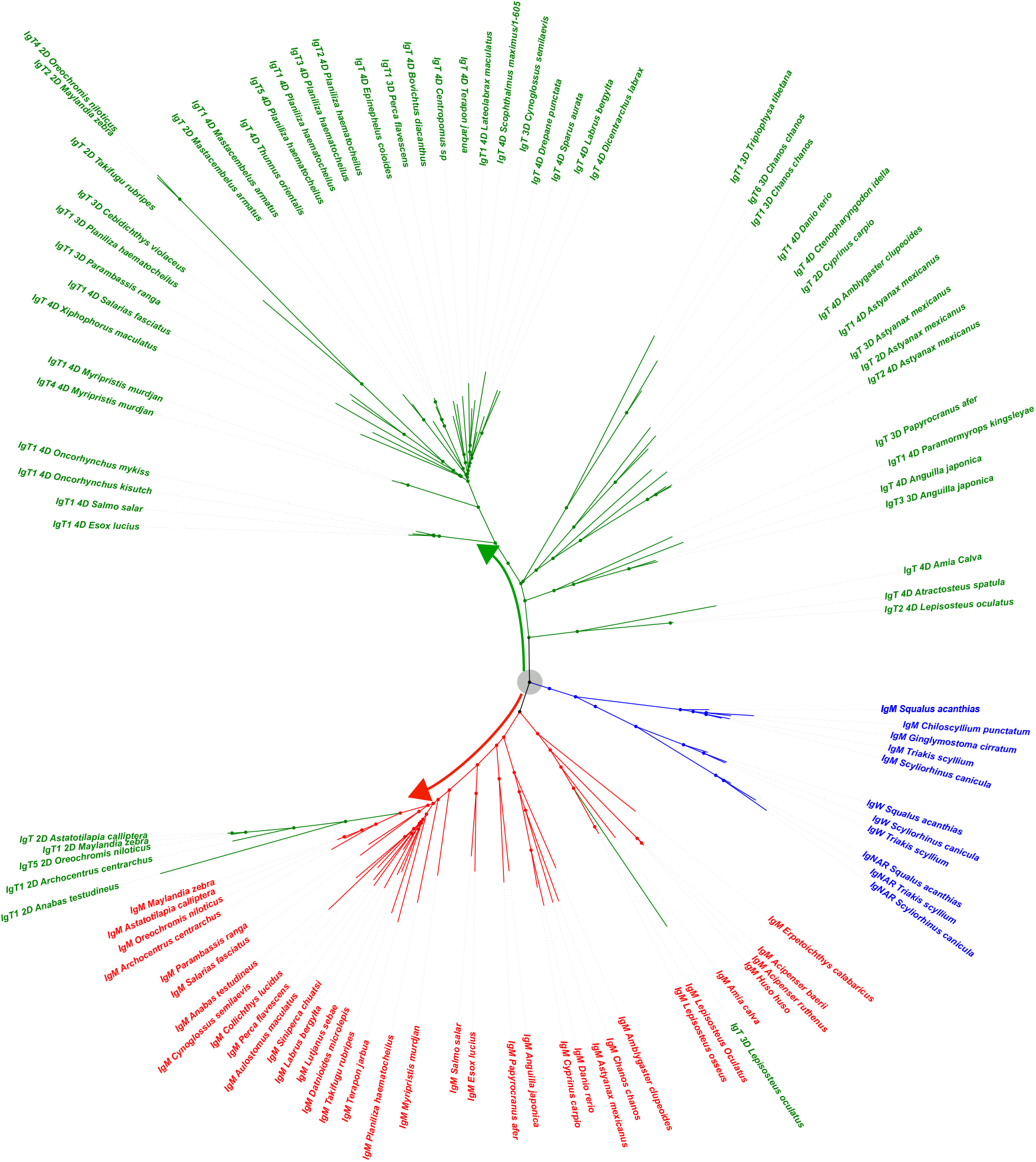
The phylogenetic tree consisting of 105 immunoglobulin sequences. The identified clades are as follows: IgT of Neopterygii (green), IgM of Actinopterygii (red), and the immunoglobulins of Elasmobranchii (blue). IgT 4D, IgT 3D, and IgT 2D refer to the number of immunoglobulin domains that IgT exhibits. Gray circles mark the divergence points of immunoglobulins, taking as reference their appearance in Elasmobranchii.

A study of the IgT domains in holosteos is of particular interest, since until now, the oldest IgTs described in the scientific literature correspond to those of Cypriniformes. In addition, these observations of IgT in other teleost species whose origin is prior to that of Cypriniformes, have not been previously described. As such, the large number of these IgT sequences provides an opportunity to study the evolutionary events that may underlie the origin of IgT (Figure 4). In 53 IgT sequences, exon coding CH1T is very similar to that of CH1M (taxa are in the same clade), while 80 other sequences are within a clade related to CH1M. These results suggest that the exon coding CH1 of the Neopterygii IgT shows a tendency to be exchanged for the exon coding CH1 of the IgM of the same species; in some cases, this occurred quite recently, since the CH1M and CH1T sequences have similarities of more than 98%. Such phenomena is already observed early in the IgT3D of *Lepisosteus oculatus*.

**Figure 4:**
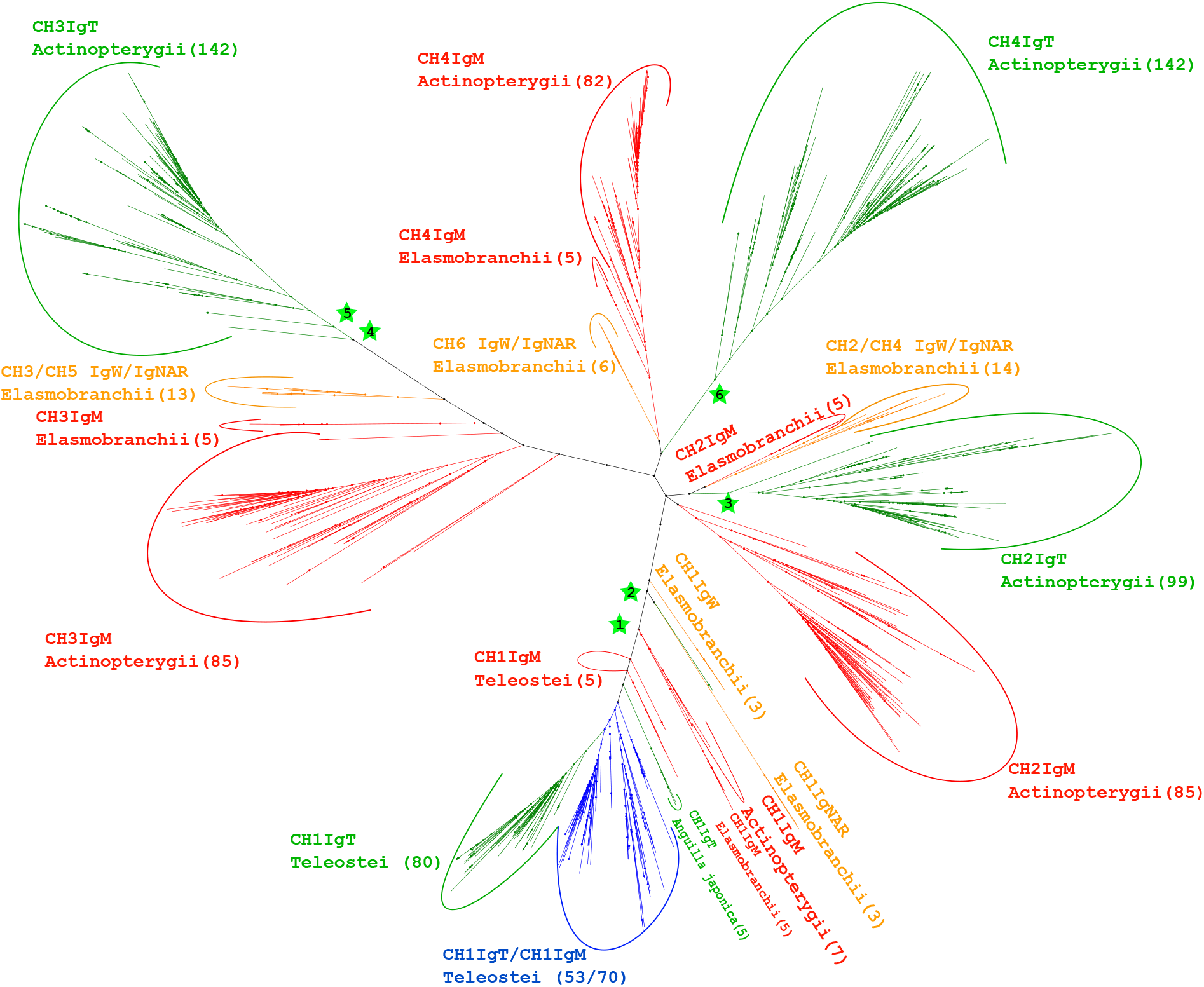
The phylogenetic tree of the domain study of the 252 immunoglobulin sequences used in this work. The number of domains per clade is provided in brackets. The immunoglobulin domains are labelled as follows: IgM of Actinopterygii and Elasmobranchii (red), the Neopterygii IgT (green), those of IgW and IgNAR of Elasmobranchii (orange), and a mixed clade with IgM and IgT CH1 domains (blue). Asterisks are used to indicate the presence of a domain for the IgT of holosteos. In particular, these are as follows: (asterisk 1) the CH1 IgT of the IgT3D of *Lepisosteus oculatus*, (asterisk 2) the CH1 IgT domain of the Holosteos IgT residue, (asterisk 3) the CH2 IgT from Holosteos, (asterisk 4) the CH3 IgT of IgT3D, (asterisk 5) the CH3 IgT domain of the Holosteos IgT residue, and (asterisk 6) the CH4 IgT from Holosteos.

The CH2 and CH3 domains of IgT show a greater evolutionary proximity to CH2 and CH3 of IgW and IgNAR than to those of Actinopterygii IgM. Finally, the CH4 of IgT generates a clade that is evolutionary distant from the rest of the immunoglobulins. This may be because its origin coincides with the rise of immunoglobulins, or because of rapid evolutionary processes of this domain driven by selective pressure.

### 3.3. IgT absence in Teleostei

From the analysis with CHfinder, genes were not found for the IgT in 25 species out of the 73 Actinopterygii studied (see Table S1). The absence in two of these species (*Epinephelus lanceolatus* and *Datnioides undecimradiatus*) can be explained by assemble problems (presence of gaps in the locus that contain the immunoglobulins). In other cases, one species of Siluriform (channel catfish) and another of Beloniformes (medaka) had already been described as lacking the gene for the IgT (Magadán-Mompó et al., 2011; Bengtén et al., 2006). Here, we also did not find this gene in three other Siluriformes species as well as another from the Beloniformes. In all other fish species studied, the loss of IgT is new information that has not been previously described. For a complete listing, refer to Table S4 and Figure 5.

**Figure 5:**
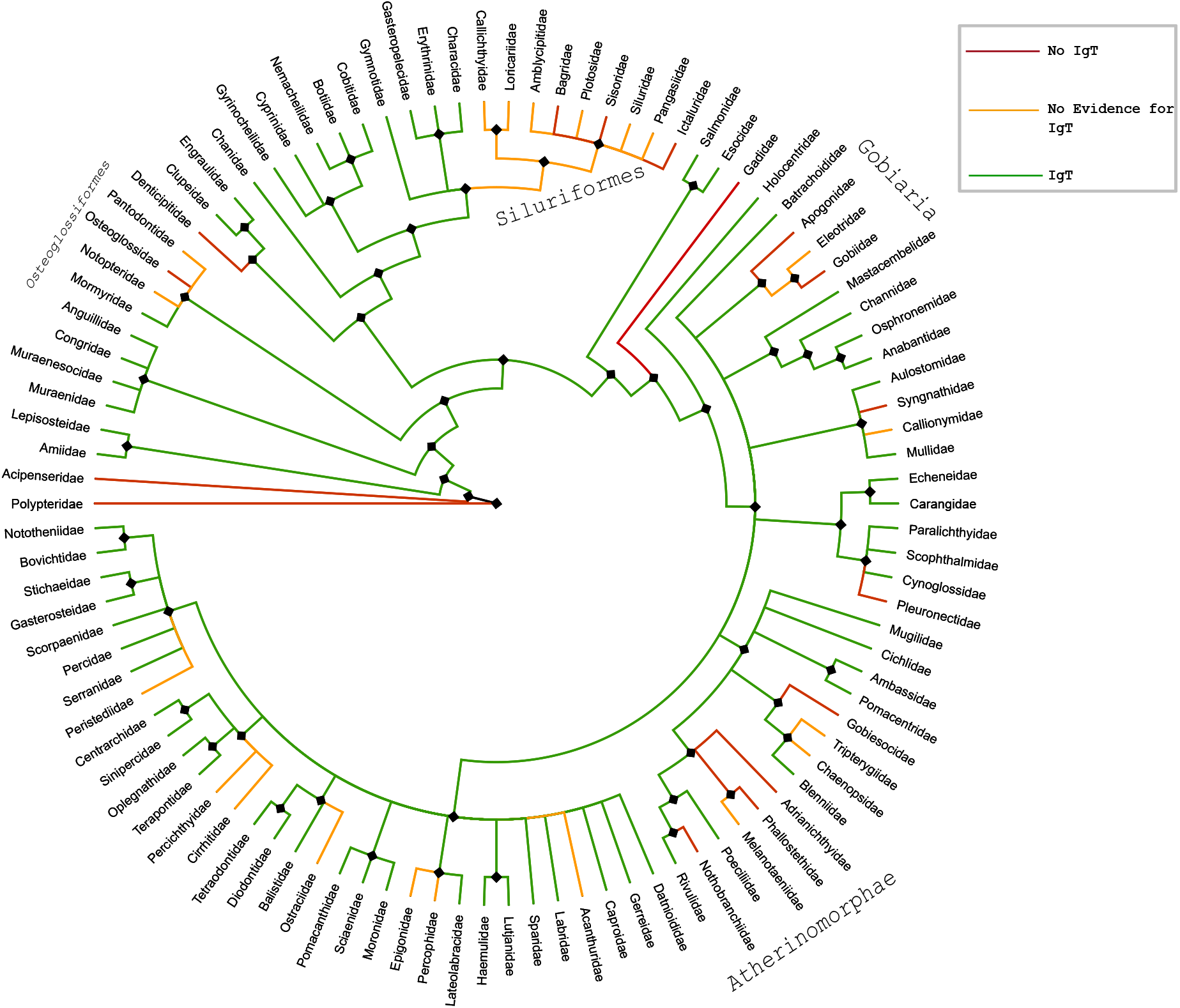
Cladogram from the data obtained in Table **??**. Clades in green indicate the presence of IgT in at least one member of that family. The red clades indicate the loss of IgT in the members analyzed for that family. Those clades with insufficient data to determine the absence of IgT are colored orange.

## 4. Discussion

At present, there is an increasing availability of high-quality genomes and transcriptomes of fish species. By using machine learning, the CHfinder program has allowed us to study fish Igs in a fast and agile manner. The program identifies the majority of the CH exons in available genomes with high accuracy. The absolute number of CH exons may be an underestimate in some cases due to the early stages of sequencing projects and the presence of gaps. For genome searches, it does not attempt to differentiate whether an exon sequence belongs to a pseudogene or a viable gene. The program provides information about the presence of homologous exon sequences to high precision. As such, it is valid for the identification of genes and obtaining sequences for evolutionary studies.

Previous publications suggest that IgT had its origin through duplication of the IgM gene.

However, this work indicate a more complex origin. CH1T is evolutionarily related to CH1M and there are numerous species with identical sequences in these two domains. Probably the need to receive the VDJ complex is the basis of its similarities and the cause of its integration into IgT. The CH2T and CH3T domains by evolutionary proximity are closer to the CH2 and CH3 domains of IgW and IgNAR of Elasmobranchii than to those of IgM. Finally, the CH4 domain raises doubts as to its origin due to the high divergence that it presents against its IgM and of the IgW and IgNAR counterparts, suggesting the possibility that this domain is so ancient that it directly connects with the appearance of the immunoglobulins themselves. In 2016, the genome of *Lepisosteus oculatus* was sequenced and the presence of two Immunoglobulin loci was noted (Braasch et al., 2016). In this work, we describe two new IgTs, one with 4 domains located in all Holostei fishes and one with 3 domains in spotted gar (IgT3D). Holostei have approximately 320 million years of independent evolution with respect to Teleostei. The CH2 and CH3 domains of the Holostei IgT are more similar to IgW domains, suggestive of their probable origin. IgT3D presents two peculiarities: it does not present the CH2 domain and the CH1 domain has a high similarity with the CH1 of the IgM, indicating a recent exchange. Both observations are in accordance with previous works in Teleosts. The presence of IgT in Holostei show that it is not a specific immunoglobulin of Teleostei.

Here, we found that both Holostei and Telostei possess IgT, contrary to the belief that it is a specific teleost immunoglobulin. Its origin would be approximately 300 Mya, coinciding with the late Permian and the appearance of the Neopterygii. This is supported by the fact that we have not found IgT in Acipenseriformes or in Polypteriformes. However, the Ig domain tree shows that CH2, CH3, and CH4 of the IgT in Neopterygii diverged prior to the occurrence of Actinopterygii, suggesting a recent loss of IgT in Acipenseriformes and Polypteriformes. This loss in fish of what seem to be key elements of the adaptive immune systems, without seeming to condition the viability of these species, is seen in several species. Examples include the loss of MHC-II in the Atlantic cod, pipefish y anglerfish (Star et al., 2011; Haase et al., 2013; Dubin et al., 2019), the loss of IgT in the medaka and channel catfish, or the loss of immunoglobulin genes in *Gouania willdenowii*.

Our work corroborates previous studies that IgT exists throughout the Neopterygii clade. Until now however, IgT loss has been documented in only two species of Teleostei. Here we describe that the extent of this loss is greater than previously observed. Moreover, this appears to be at the level of order or family. Nonetheless, the causes and consequences of the loss of this Ig isotype remains unclear. In this work we have sought to clarify the origin of IgT and described its presence in Holostei. However, several questions remain, including whether IgT arose as an IgM-IgW/NAR chimera with the emergence of the Neopterygii.

## Supplementary Material

**Table. S1:** Results of gene/exon identification with CHfinder

| Specie | IGT | IGM | IGD | Order | Family |
| --- | --- | --- | --- | --- | --- |
| **CLADISTIA** |  |  |  |  |  |
| *Erpetoichthys calabaricus* | 0 | 1 | 1 | Polypteriformes | Polypteridae |
| **ACTINOPTERI** |  |  |  |  |  |
| ->CHONDROSTEI |  |  |  |  |  |
| *Acipenser ruthenus* | 0 | 2 | 3 | Acipenseriformes | Acipenseridae |
| ->NEOPTERYGII |  |  |  |  |  |
| ->HOLOSTEI |  |  |  |  |  |
| *Lepisosteus oculatus* | 2 | 1 | 1 | Lepisosteiformes | Lepisosteidae |
| ->TELEOSTEI |  |  |  |  |  |
| —>ELOPOCEPHALA |  |  |  |  |  |
| *Anguilla japonica* | 5 | 1 | 1 | Anguilliformes | Anguillidae |
| —>OSTEOGLOSSOCEPHALA |  |  |  |  |  |
| *Scleropages formosus* | 0 | 2 | 2 | Osteoglossiformes | Osteoglossidae |
| *Paramormyrops kingsleyae* | 2 | 1 | 1 | Osteoglossiformes | Mormyridae |
| —>CLUPEOCEPHALA |  |  |  |  |  |
| —>OTOMORPHA |  |  |  |  |  |
| *Clupea harengus* | 1 | 2 | 2 | Clupeiformes | Clupeidae |
| *Coilia nasus* | 2 | 1 | 1 | Clupeiformes | Engraulidae |
| *Denticeps clupeoides* | 0 | 1 | 1 | Clupeiformes | Denticipitidae |
| *Chanos chanos* | 6 | 2 | 2 | Gonorynchiformes | Chanidae |
| *Carassius auratus* | 4 | 2 | 2 | Cypriniformes | Cyprinidae |
| *Danio rerio* | 1 | 1 | 1 | Cypriniformes | Cyprinidae |
| *Cyprinus carpio* | 2 | 2 | 3 | Cypriniformes | Cyprinidae |
| *Triplophysa tibetana* | 2 | 2 | 2 | Cypriniformes | Nemacheilidae |
| *Electrophorus electricus* | 1 | 1 | 1 | Gymnotiformes | Gymnotidae |
| *Bagarius yarrelli* | 0 | 1 | 1 | Siluriformes | Sisoridae |
| *Ictalurus punctatus* | 0 | 1 | 2 | Siluriformes | Ictaluridae |
| *Tachysurus fulvidraco* | 0 | 1 | 1 | Siluriformes | Bagridae |
| *Pangasianodon hypophthalmus* | 0 | Gap | 1 | Siluriformes | Pangasiidae |
| *Astyanax mexicanus* | 5 | 1 | 3 | Characiformes | Characidae |
| —>EUTELEOSTEOMORPHA |  |  |  |  |  |
| —>PROTACANTHOPTERYGII |  |  |  |  |  |
| *Salmo trutta* | 3 | 2 | 2 | Salmoniformes | Salmonidae |
| *Oncorhynchus mykiss* | 3 | 2 | 4 | Salmoniformes | Salmonidae |
| *Oncorhynchus tshawytscha* | 1 | 2 | 2 | Salmoniformes | Salmonidae |
| *Oncorhynchus kisutch* | 3 | 1 | 2 | Salmoniformes | Salmonidae |
| *Coregonus* | 3 | 1 | 3 | Salmoniformes | Salmonidae |
| *Esox lucius* | 2 | 2 | 2 | Esociformes | Esocidae |
| ——>NEOTELEOSTEI |  |  |  |  |  |
| ——>PARACANTHOMORPHACEA |  |  |  |  |  |
| *Gadus morhua* | 0 | 3 | 3 | Gadiformes | Gadidae |
| ——>HOLOCENTRIMORPHACEAE |  |  |  |  |  |
| *Myripristis murdjan* | 5 | 8 | 8 | Holocentriformes | Holocentridae |
| ——>PERCOMORPHACEAE |  |  |  |  |  |
| ——>BATRACHOIDIARIA |  |  |  |  |  |
| *Thalassophryne amazonica* | 1 | 2 | 0 | Batrachoidiformes | Batrachoididae |
| ——>PERCOMORPHACEAE |  |  |  |  |  |
| ——>EUPERCARIA |  |  |  |  |  |
| *Labrus bergylta* | 7 | 12 | 16 | Labriformes | Labridae |
| *Notolabrus celidotus* | 2 | 1 | 1 | Labriformes | Labridae |
| *Takifugu rubripes* | 1 | 1 | 1 | Tetraodontiformes | Tetraodontidae |
| *Larimichthys crocea* | 1 | 2 | 1 | Acanthuriformes | Sciaenidae |
| *Sparus aurata* | 5 | 2 | 2 | Spariformes | Sparidae |
| *Cebidichthys violaceus* | 1 | 2 | 2 | Perciformes | Stichaeidae |
| *Cottoperca gobio* | 1 | 1 | 1 | Perciformes | Bovichtidae |
| *Perca flavescens* | 3 | 1 | 1 | Perciformes | Percidae |
| *Plectropomus leopardus* | 1 | 1 | 1 | Perciformes | Serranidae |
| *Epinephelus lanceolatus* | Gap | 2 | 1 | Perciformes | Serranidae |
| *Epinephelus moara* | 0 | 1 | 1 | Perciformes | Serranidae |
| *Oplegnathus fasciatus* | 2 | 1 | 1 | Centrarchiformes | Oplegnathidae |
| *Lateolabrax maculatus* | 2 | 2 | 2 | Pempheriformes | Lateolabracidae |
| *Collichthys lucidus* | 1 | 4 | 3 | I. s. in Eupercaria | Sciaenidae |
| *Datnioides undecimradiatus* | Gap | 1 | 1 | Lobotiformes | Datnioididae |
| ——>PERCOMORPHACEAE |  |  |  |  |  |
| ——>GOBIARIA |  |  |  |  |  |
| *Periophthalmus magnuspinnatus* | 0 | 1 | 1 | Gobiiformes | Gobiidae |
| *Neogobius melanostomus* | 0 | 2 | 2 | Gobiiformes | Gobiidae |
| *Sphaeramia orbicularis* | 0 | 1 | 2 | Kurtiformes | Apogonidae |
| ——>PERCOMORPHACEAE |  |  |  |  |  |
| ——>SYNGNATHARIA |  |  |  |  |  |
| *Syngnathus acus* | 0 | 1 | 1 | Syngnathiformes | Syngnathidae |
| ——>PERCOMORPHACEAE |  |  |  |  |  |
| ——>ANABANTARIA |  |  |  |  |  |
| *Anabas testudineus* | 3 | 1 | 3 | Anabantiformes | Anabantidae |
| *Betta splendens* | 0 | 1 | 1 | Anabantiformes | Osphronemidae |
| *Channa argus* | 1 | 1 | 4 | Anabantiformes | Channidae |
| *Mastacembelus armatus* | 4 | 1 | 1 | Synbranchiformes | Mastacembelidae |
| ——>PERCOMORPHACEAE |  |  |  |  |  |
| ——>CARANGARIA |  |  |  |  |  |
| *Cynoglossus semilaevis* | 1 | 1 | 1 | Pleuronectiformes | Cynoglossidae |
| *Scophthalmus maximus* | 1 | 1 | 1 | Pleuronectiformes | Scophthalmidae |
| *Reinhardtius hippoglossoides* | 0 | 1 | 1 | Pleuronectiformes | Pleuronectidae |
| *Hippoglossus hippoglossus* | 0 | 1 | 1 | Pleuronectiformes | Pleuronectidae |
| *Echeneis naucratis* | 1 | 3 | 3 | Carangiiformes | Echeneidae |
| ——>PERCOMORPHACEAE |  |  |  |  |  |
| ——>OVALENTARIA |  |  |  |  |  |
| *Planiliza haematocheilus* | 20 | 4 | 4 | Mugiliformes | Mugilidae |
| *Archocentrus centrarchus* | 4 | 4 | 4 | Cichliiformes | Cichlidae |
| *Oreochromis niloticus* | 5 | 6 | 6 | Cichliiformes | Cichlidae |
| *Astatotilapia calliptera* | 1 | 2 | 3 | Cichliiformes | Cichlidae |
| *Maylandia zebra* | 4 | 2 | 3 | Cichliiformes | Cichlidae |
| *Neostethus bicornis* | 0 | 1 | 1 | Atheriniiformes | Phallostethidae |
| *Poecilia reticulata* | 2 | 1 | 1 | Cyprinodontiformes | Poeciliidae |
| *Xiphophorus maculatus* | 2 | 1 | 1 | Cyprinodontiformes | Poeciliidae |
| *Nothobranchius furzeri* | 0 | 1 | 2 | Cyprinodontiformes | Nothobranchiidae |
| *Kryptolebias hermaphroditus* | 2 | 2 | 2 | Cyprinodontiformes | Rivulidae |
| *Oryzias latipes* | 0 | 6 | 6 | Beloniiformes | Adrianichthyidae |
| *Oryzias javanicus* | 0 | 3 | 3 | Beloniiformes | Adrianichthyidae |
| *Salarias fasciatus* | 5 | 9 | 11 | Blenniiformes | Blenniidae |
| *Gouania willdenowi* | 0 | 0 | 0 | Blenniiformes | Gobiesocidae |
| *Parambassis ranga* | 4 | 1 | 2 | I. s. in Ovalentaria | Ambassidae |
| *Amphiprion percula* | 0 | 4 | 6 | I. s. in Ovalentaria | Pomacentridae |

**Figure S1:**
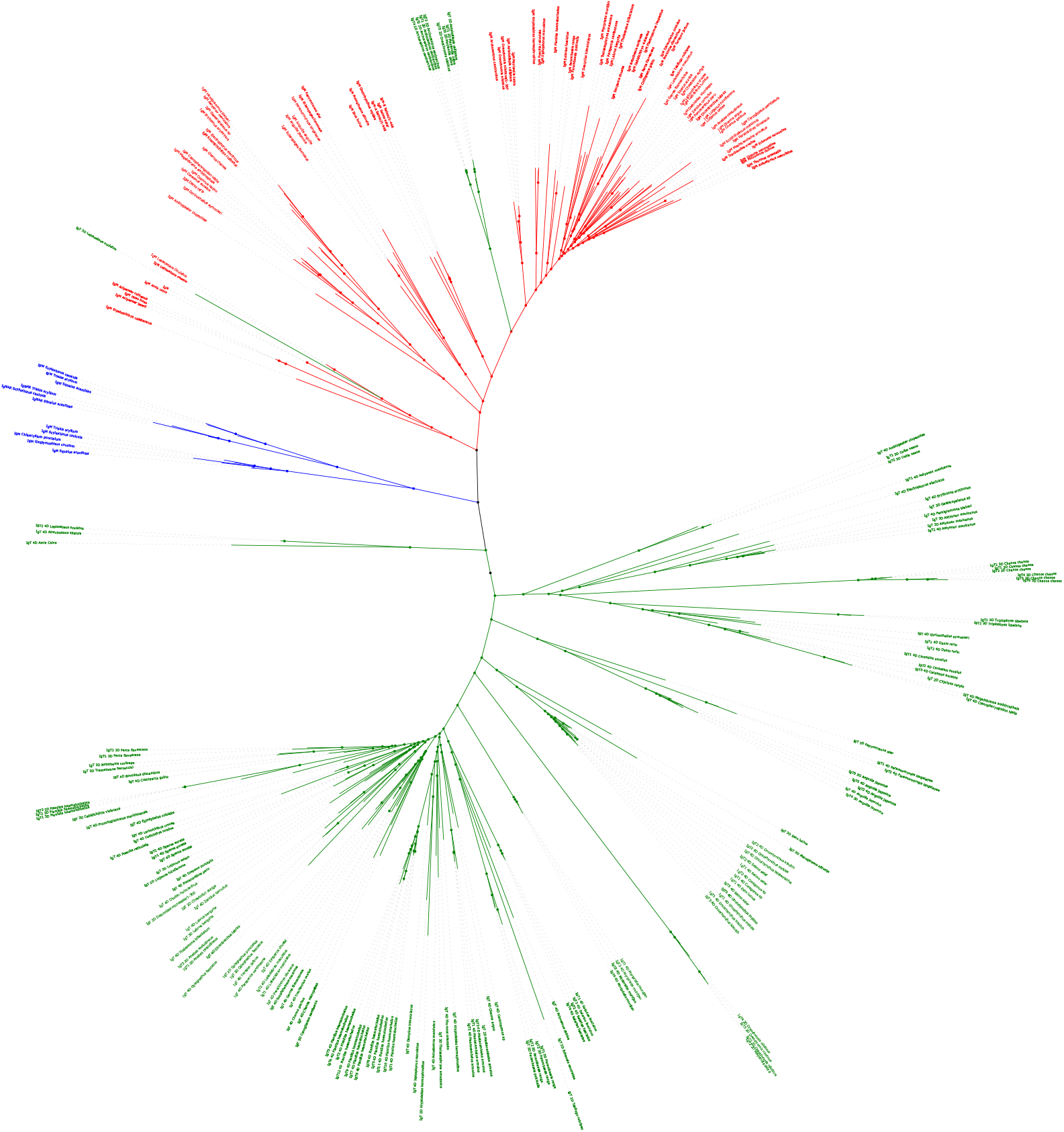
The phylogenetic tree that includes the 252 immunoglobulin sequences used in this work. The clades are labelled as follows: IgT of Neopterygii (green), the IgM of Actinopterygii (red), Ig of Elasmobranchii (blue). The labels: IgT 4D, IgT 3D, and IgT 2D refer to the number of immunoglobulin domains that IgT exhibits.

**Table. S4:** Data for the IgT cladogram.

| Order | Family | Genome | Fish T1K | Bibliography |
| --- | --- | --- | --- | --- |
| Polypteriformes (3) | Polypteridae (3) | NO (1) | NO (3) | NO |
| Acipenseriformes (2) | Acipenseridae<br>Polyodontidae | NO (1)<br>No Genome | No sample<br>NO (1) | NO<br>NO |
| Lepisosteiformes (3) | Lepisosteidae | NO (1) | NO (2) | NO |
| Anguilliformes (4) | Anguillidae<br>Muraenidae<br>Muraenesocidae<br>Congridae | YES (1)<br>No Genome<br>No Genome<br>No Genome | YES (1)<br>NO (1)<br>NO (1)<br>NO (1) | NO<br>NO<br>NO<br>NO |
| Osteoglossiformes (5) | Osteoglossidae<br>Notopteridae<br>Pantodontidae<br>Mormyridae | NO (1)<br>No Genome<br>No Genome<br>YES (1) | NO (1)<br>NO (1)<br>NO (1)<br>YES (1) | NO<br>NO<br>NO<br>NO |
| Clupeiformes (4) | Clupeidae<br>Engraulidae<br>Denticipitidae | YES (1)<br>YES (1)<br>NO (1) | YES (1)<br>No sample<br>No sample | NO<br>NO<br>NO |
| Gonorynchiformes (1) | Chanidae | YES (1) | No sample | NO |
| Cypriniformes (13) | Cyprinidae<br>Nemacheilidae<br>Botiidae<br>Gyrinocheilidae<br>Cobitidae | YES (3)<br>YES (1)<br>No Genome<br>No Genome<br>No Genome | YES (1)<br>NO (1)<br>YES (1)<br>YES (1)<br>YES (1) | YES (5)<br>NO<br>NO<br>NO<br>YES (1) |
| Gymnotiformes (2) | Gymnotidae | YES (1) | YES (2) | No |
| Siluriformes (11) | Sisoridae<br>Ictaluridae<br>Bagridae<br>Pangasiidae<br>Siluridae<br>Loricariidae<br>Plotosidae<br>Amblycipitidae | NO (1)<br>NO (1)<br>NO (1)<br>NO (1)<br>No Genome<br>No Genome<br>No Genome<br>No Genome | NO (1)<br>NO (1)<br>NO (2)<br>No sample<br>NO (1)<br>NO (1)<br>NO (1)<br>NO (1) | NO<br>NO<br>NO<br>NO<br>NO<br>NO<br>NO<br>NO |
|  | Callichthyidae | No Genome | NO (1) | NO |
| Characiformes (4) | Characidae<br>Gasteropelecidae<br>Erythrinidae | YES (1)<br>No Genome<br>No Genome | YES (1)<br>YES (1)<br>YES (1) | NO<br>NO<br>NO |
| Salmoniformes (6) | Salmonidae | YES (5) | No sample | YES(2) |
| Esociformes (1) | Esocidae | YES (1) | No sample | NO |
| Gadiformes (2) | Gadidae | NO (1) | NO (1) | NO |
| Holocentriformes (4) | Holocentridae | YES (1) | YES (3) | NO |
| Batrachoidiformes(2) | Batrachoididae | YES (1) | YES (1) | NO |
| Labriformes (4) | Labridae | YES (2) | YES (2) | YES (1) |
| Tetraodontiformes (5) | Tetraodontidae<br>Balistidae<br>Diodontidae<br>Ostraciidae | YES (1)<br>No Genome<br>No Genome<br>No Genome | No sample<br>YES (1)<br>YES (1)<br>NO (2) | YES (1)<br>NO<br>NO<br>NO |
| Acanthuriformes (3) | Sciaenidae<br>Zanclidae<br>Acanthuridae | YES (1)<br>No Genome<br>No Genome | YES (1)<br>NO (1) | NO<br>NO<br>NO |
| Spariformes (1) | Sparidae | YES (1) | No sample | YES (1) |
| Perciformes (16) | Stichaeidae<br>Bovichtidae<br>Percidae<br>Serranidae<br>Nototheniidae<br>Gasterosteidae<br>Scorpaenidae<br>Peristediidae | YES (1)<br>YES (1)<br>YES (1)<br>YES/NO (1/2)<br>No Genome<br>No Genome<br>No Genome<br>No Genome | No sample<br>No sample<br>No sample<br>YES (1)<br>No sample<br>No sample<br>YES/NO (2/2)<br>NO (1) | NO<br>YES (1)<br>NO<br>YES (1)<br>YES (2)<br>YES (1)<br>NO<br>NO |
| Centrarchiformes (8) | Oplegnathidae<br>Sinipercaidae<br>Terapontidae | YES (1)<br>No Genome<br>No Genome | YES (1)<br>YES (1)<br>YES (1) | NO<br>YES (1)<br>NO |
|  | Centrarchidae | No Genome | YES (1) | NO |
|  | Cirrhitidae | No Genome | NO (1) | NO |
|  | Percichthyidae | No Genome | NO (1) | NO |
| Pempheriformes (3) | Lateolabracidae | YES (1) | No sample | NO |
|  | Epigonidae | No Genome | NO (1) | NO |
|  | Percophidae | No Genome | NO (1) | NO |
| I. s. in Eupercaria (10) | Sciaenidae | YES (1) | YES (1) | NO |
|  | Lutjanidae | No Genome | YES (2) | NO |
|  | Pomacanthidae | No Genome | YES (1) | NO |
|  | Gerreidae | No Genome | YES (2) | NO |
|  | Haemulidae | No Genome | YES (1) | NO |
|  | Caproidae | No Genome | YES (1) | NO |
|  | Moronidae | No Genome | No sample | YES (1) |
| Lobotiformes (2) | Datnioididae | NO (1) | YES (1) | NO |
| Gobiiformes (5) | Gobiidae | NO (2) | NO (2) | NO |
|  | Eleotridae | No Genome | NO (1) | NO |
| Kurtiformes (3) | Apogonidae | NO (1) | NO (2) | NO |
| Syngnathiformes (2) | Syngnathidae | NO (1) | NO (1) | NO |
|  | Mullidae | No Genome | YES (1) | NO |
|  | Callionymidae | No Genome | NO (1) | NO |
|  | Aulostomidae | No Genome | YES (1) | NO |
| Anabantiformes (6) | Anabantidae | YES (1) | NO (1) | NO |
|  | Osphronemidae | NO (1) | YES (1) | NO |
|  | Channidae | YES (1) | YES (2) | NO |
| Synbranchiformes (2) | Mastacembelidae | YES (1) | NO (2) | NO |
| Pleuronectiformes (5) | Cynoglossidae | YES (1) | No sample | NO |
|  | Scophthalmidae | NO (1) | NO (1) | YES (1) |
|  | Pleuronectidae | NO (2) | No sample | NO |
|  | Paralichthyidae | No Genome | No sample | YES (1) |
| Carangiformes (3) | Echeneidae | YES (1) | No sample | NO |
|  | Carangidae | No Genome | YES (2) | NO |
| Mugiliformes | Mugilidae | YES (1) | YES/NO (1/1) | NO |
| Cichliformes (6) | Cichlidae | YES (4) | NO (2) | YES |
| Atheriniformes | Phallostethidae<br>Melanotaeniidae | NO (1)<br>No Genome | No sample<br>NO (2) | NO<br>NO |
| Cyprinodontiformes | Poeciliidae<br>Nothobranchiidae<br>Rivulidae | YES (2)<br>NO (1)<br>YES | YES (2)<br>No sample<br>No sample | NO<br>NO<br>NO |
| Beloniformes (3) | Adrianichthyidae | NO (2) | NO (1) | NO |
| Blenniiformes | Blenniidae<br>Gobiesocidae<br>Tripterygiidae<br>Chaenopsidae | YES (1)<br>NO (1)<br>No Genome<br>No Genome | No sample<br>NO (2)<br>NO (1)<br>NO (1) | NO<br>NO<br>NO<br>NO |
| I. s. in Ovalentaria (5) | Ambassidae<br>Pomacentridae | YES (1)<br>NO (1) | (1)<br>YES (2) | NO<br>NO |

**Table. S2:** List of fishes with IgT and representative bibliographic references.

| IgT of different Teleostei published in scientific articles |  |  |  |  |  |  |
| --- | --- | --- | --- | --- | --- | --- |
| Specie | IGT | Domain | Order | Family | Protein NCBI | Bibliography |
| *Megalabrama amblycephala* | 1 | 4 | Cypriniformes | Cyprinidae | AGR34024.1 | (Xia et al., 2016) |
| *Ctenopharyngodon idella* | 1 | 4 | Cypriniformes | Cyprinidae | ADD82655.1 | (Xiao et al., 2010) |
| *Cyprinus carpio* | 1 | 2(1,4) | Cypriniformes | Cyprinidae | BAD69715.1 | (Savan et al., 2005a) |
| *Cyprinus carpio* | 1 | 4 | Cypriniformes | Cyprinidae | BAJ41037.1 | (Ryo et al., 2010) |
| *Danio rerio* | 1 | 4 | Cypriniformes | Cyprinidae | AAT67444.1 | (Danilova et al., 2005) |
| *Danio rerio* | 1 | 4 | Cypriniformes | Cyprinidae | ACH92959.1 | (Hu et al., 2010) |
| *Oncorhynchus mykiss* | 3 | 4 | Salmoniformes | Salmonidae | AAW66978.1 | (Zhang et al., 2017) |
| *Oncorhynchus mykiss* | 3 | 4 | Salmoniformes | Salmonidae | AAV48553.1 | (Zhang et al., 2017) |
| *Oncorhynchus mykiss* | 3 | 4 | Salmoniformes | Salmonidae | ANW11927.1 | (Zhang et al., 2017) |
| *Salmo salar* | 3 | 4 | Salmoniformes | Salmonidae | ADD59873.1 | (Yasuike et al., 2010) |
| *Salmo salar* | 3 | 4 | Salmoniformes | Salmonidae | ADD59858.1 | (Yasuike et al., 2010) |
| *Salmo salar* | 3 | 4 | Salmoniformes | Salmonidae | ADD59859.1 | (Yasuike et al., 2010) |
| *Plecoglossus altivelis* | 1 | 3(1,2,4) | Osmeriformes | Plecoglossidae | BAP75403.1 | (Kato et al., 2015) |
| *Takifugu rubripes* | 1 | 2(1,4) | Tetraodontiformes | Tetraodontidae | BAD69712.1 | (Savan et al., 2005b) |
| *Sparus aurata* | 1 | 4 | Spariformes | Sparidae | ASK39431.1 | (Piazzon et al., 2016) |
| *Bovichtus diacanthus* | 1 | 4 | Perciformes | Bovichtidae | AKA09828.1 | (Giacomelli et al., 2015) |
| *Trematomus bernacchii* | 3 | 3(1,3,4) | Perciformes | Nototheniidae | AKA09825.1 | (Giacomelli et al., 2015) |
| *Notothenia coriiceps* | ? | 3(1,3,4) | Perciformes | Nototheniidae | AKA09827.1 | (Giacomelli et al., 2015) |
| *Epinephelus coioides* | 1 | 4 | Perciformes | Serranidae | ACZ54909.1 | (Du et al., 2016) |
| *Siniperca chuatsi* | 1 | 4 | Centrarchiformes | Sinipercidae | AAY42141 | (Tian et al., 2009) |
| *Thunnus orientalis* | 1 | 4 | Scombriformes | Scombridae | AHC31432 | (Mashoof et al., 2014) |
| *Dicentrarchus labrax* | 1 | 4 | Eupercaria I.S. | Moronidae | AKK32388.1 | (Buonocore et al., 2017) |
| *Scophthalmus maximus* | 1 | 4 | Pleuronectiformes | Scophthalmidae | AMQ49169.1 | (Tang et al., 2018) |
| *Paralichthys olivaceus* | 1 | 4 | Pleuronectiformes | Paralichthyidae | ANS12794 | (Du et al., 2016) |
| *Oreochromis niloticus* | 1 | 2(1,4) | Cichliformes | Cichlidae | AKN35078 | (Velázquez et al., 2018) |
| *Labeo catla* | 1 | ? | Cypriniformes | Cyprinidae | ALF45197.1 | (Patel et al., 2016) |
| *Labeo rohita* | ? | ? | Cypriniformes | Cyprinidae | ————— | (Kar et al., 2015) |
| *Misgurnus anguillicaudatus* | 1 | 4 | Cypriniformes | Cobitidae | ————— | (Xu et al., 2019) |
| *Gasterosteus aculeatus* | 1 | 3(1,3,4) | Perciformes | Gasterosteidae | ————— | (Gambón-Deza et al., 2010) |
| *Labrus bergylta* | 1 | 4 | Labriformes | Labridae | ————— | (Bilal et al., 2019) |

**Table S3:**
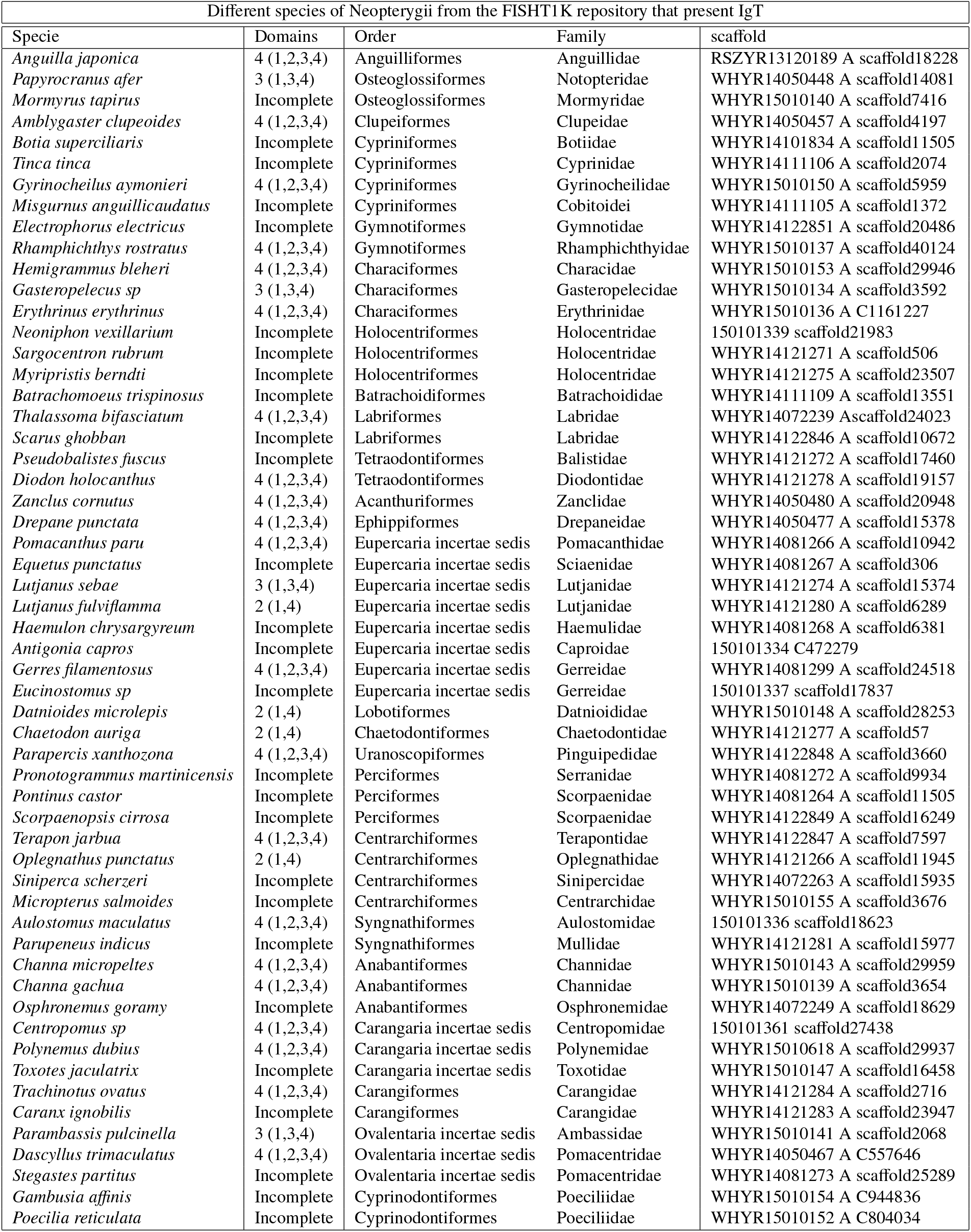
IgT in Fish T1K

## References

Afgan, E., Baker, D., Batut, B., Van Den Beek, M., Bouvier, D., Čech, M., Chilton, J., Clements, D., Coraor, N., Grüning, B. A. et al. (2018). The galaxy platform for accessible, reproducible and collaborative biomedical analyses: 2018 update. Nucleic acids research, 46, W537–W544.

Bengtén, E., Clem, L. W., Miller, N. W., Warr, G. W., & Wilson, M. (2006). Channel catfish immunoglobulins: repertoire and expression. Developmental & Comparative Immunology, 30, 77–92.

Bengtén, E., Leanderson, T., & Pilström, L. (1991). Immunoglobulin heavy chain cDNA from the teleost Atlantic cod (Gadus morhua l.): nucleotide sequences of secretory and membrane form show an unusual splicing pattern. European journal of immunology, 21, 3027–3033.

Bengtén, E., Quiniou, S. M.-A., Stuge, T. B., Katagiri, T., Miller, N. W., Clem, L. W., Warr, G. W., & Wilson, M. (2002). The IgH locus of the channel catfish, Ictalurus punctatus, contains multiple constant region gene sequences: different genes encode heavy chains of membrane and secreted IgD. The Journal of Immunology, 169, 2488–2497.

Bilal, S., Lie, K. K., Dalum, A. S., Karlsen, O. A., & Hordvik, I. (2019). Analysis of immunoglobulin and T cell receptor gene expression in ballan wrasse (labrus bergylta) revealed an extraordinarily high IgM expression in the gut. Fish & shellfish immunology, 87, 650–658.

Braasch, I., Gehrke, A. R., Smith, J. J., Kawasaki, K., Manousaki, T., Pasquier, J., Amores, A., Desvignes, T., Batzel, P., Catchen, J. et al. (2016). The spotted gar genome illuminates vertebrate evolution and facilitates human-teleost comparisons. Nature genetics, 48, 427–437.

Buonocore, F., Stocchi, V., Nunez-Ortiz, N., Randelli, E., Gerdol, M., Pallavicini, A., Facchiano, A., Bernini, C., Guerra, L., Scapigliati, G. et al. (2017). Immunoglobulin T from sea bass (Dicentrarchus labrax L.): molecular characterization, tissue localization and expression after nodavirus infection. BMC molecular biology, 18, 8.

Castro, C. D., & Flajnik, M. F. (2014). Putting J chain back on the map: how might its expression define plasma cell development? The Journal of Immunology, 193, 3248–3255.

Chen, K., Xu, W., Wilson, M., He, B., Miller, N. W., Bengten, E., Edholm, E.-S., Santini, P. A., Rath, P., Chiu, A. et al. (2009). Immunoglobulind enhances immune surveillance by activating antimicrobial, proinflammatory and B cell–stimulating programs in basophils. Nature immunology, 10, 889.

Cooper, M. D., & Alder, M. N. (2006). The evolution of adaptive immune systems. Cell, 124, 815–822.

Danilova, N., Bussmann, J., Jekosch, K., & Steiner, L. A. (2005). The immunoglobulin heavy-chain locus in zebrafish: identification and expression of a previously unknown isotype, immunoglobulin Z. Nature immunology, 6, 295.

Du, Y., Tang, X., Zhan, W., Xing, J., & Sheng, X. (2016). Immunoglobulin tau heavy chain (IgT) in flounder, Paralichthys olivaceus: molecular cloning, characterization, and expression analyses. International journal of molecular sciences, 17, 1571.

Dubin, A., Jørgensen, T. E., Moum, T., Johansen, S. D., & Jakt, L. M. (2019). Complete loss of the MHC II pathway in an anglerfish, Lophius piscatorius. Biology letters, 15, 20190594.

Edelman, G. M., Cunningham, B. A., Gall, W. E., Gottlieb, P. D., Rutishauser, U., & Waxdal, M. J. (1969). The covalent structure of an entire *γ*G immunoglobulin molecule. Proceedings of the National Academy of Sciences, 63, 78–85.

Flajnik, M. F., & Kasahara, M. (2010). Origin and evolution of the adaptive immune system: genetic events and selective pressures. Nature Reviews Genetics, 11, 47–59.

Gambon-Deza, F., Sánchez-Espinel, C., & Magadan-Mompo, S. (2009). The immunoglobulin heavy chain locus in the platypus (Ornithorhynchus anatinus). Molecular immunology, 46, 2515–2523.

Gambón-Deza, F., Sánchez-Espinel, C., & Magadán-Mompó, S. (2010). Presence of an unique IgT on the IGH locus in three-spined stickleback fish (Gasterosteus aculeatus) and the very recent generation of a repertoire of VH genes. Developmental & Comparative Immunology, 34, 114–122.

Ghaffari, S. H., & Lobb, C. (1989). Cloning and sequence analysis of channel catfish heavy chain cDNA indicate phylogenetic diversity within the IgM immunoglobulin family. The Journal of Immunology, 142, 1356–1365.

Giacomelli, S., Buonocore, F., Albanese, F., Scapigliati, G., Gerdol, M., Oreste, U., & Coscia, M. R. (2015). New insights into evolution of IgT genes coming from Antarctic teleosts. Marine genomics, 24, 55–68.

Gouy, M., Guindon, S., & Gascuel, O. (2009). SeaView version 4: a multiplatform graphical user interface for sequence alignment and phylogenetic tree building. Molecular biology and evolution, 27, 221–224.

Grabherr, M. G., Haas, B. J., Yassour, M., Levin, J. Z., Thompson, D. A., Amit, I., Adiconis, X., Fan, L., Raychowdhury, R., Zeng, Q. et al. (2011). Trinity: reconstructing a full-length transcriptome without a genome from RNA-Seq data. Nature biotechnology, 29, 644.

Haas, B. J., Papanicolaou, A., Yassour, M., Grabherr, M., Blood, P. D., Bowden, J., Couger, M. B., Eccles, D., Li, B., Lieber, M. et al. (2013). De novo transcript sequence reconstruction from RNA-seq using the Trinity platform for reference generation and analysis. Nature protocols, 8, 1494.

Haase, D., Roth, O., Kalbe, M., Schmiedeskamp, G., Scharsack, J. P., Rosenstiel, P., & Reusch, T. B. (2013). Absence of major histocompatibility complex class II mediated immunity in pipefish, Syngnathus typhle: evidence from deep transcriptome sequencing. Biology letters, 9, 20130044.

Hansen, J. D., Landis, E. D., & Phillips, R. B. (2005). Discovery of a unique Ig heavy-chain isotype (IgT) in rainbow trout: Implications for a distinctive B cell developmental pathway in teleost fish. Proceedings of the National Academy of Sciences, 102, 6919–6924.

Hikima, J.-i., Jung, T.-S., & Aoki, T. (2011). Immunoglobulin genes and their transcriptional control in teleosts. Developmental & Comparative Immunology, 35, 924–936.

Hordvik, I., Thevarajan, J., Samdal, I., Bastani, N., & Krossoy, B. (1999). Molecular cloning and phylogenetic analysis of the Atlantic salmon immunoglobulin D gene. Scandinavian journal of immunology, 50, 202–210.

Hu, Y.-L., Xiang, L.-X., & Shao, J.-Z. (2010). Identification and characterization of a novel immunoglobulin Z isotype in zebrafish: implications for a distinct b cell receptor in lower vertebrates. Molecular immunology, 47, 738–746.

Kar, B., Mohapatra, A., Mohanty, J., & Sahoo, P. K. (2015). Transcriptional changes in three immunoglobulin isotypes of rohu, labeo rohita in response to Argulus siamensis infection. Fish & shellfish immunology, 47, 28–33.

Kato, G., Takano, T., Sakai, T., Matsuyama, T., Sano, N., & Nakayasu, C. (2015). Cloning and expression analyses of a unique IgT in ayu plecoglossus altivelis. Fisheries science, 81, 29–36.

Katoh, K., & Standley, D. M. (2013). MAFFT multiple sequence alignment software version 7: improvements in performance and usability. Molecular biology and evolution, 30, 772–780.

Le, S. Q., & Gascuel, O. (2008). An improved general amino acid replacement matrix. Molecular biology and evolution, 25, 1307–1320.

Letunic, I., & Bork, P. (2019). Interactive Tree Of Life (iTOL) v4: recent updates and new developments. Nucleic acids research, 47, W256–W259.

Magadán-Mompó, S., Sánchez-Espinel, C., & Gambón-Deza, F. (2011). Immunoglobulin heavy chains in medaka (oryzias latipes). BMC evolutionary biology, 11, 165.

Mashoof, S., Pohlenz, C., Chen, P. L., Deiss, T. C., Gatlin III, D., Buentello, A., & Criscitiello, M. F. (2014). Expressed IgH *μ* and *τ* transcripts share diversity segment in ranched Thunnus orientalis. Developmental & Comparative Immunology, 43, 76–86.

Ohta, Y., & Flajnik, M. (2006). IgD, like IgM, is a primordial immunoglobulin class perpetuated in most jawed vertebrates. Proceedings of the National Academy of Sciences, 103, 10723–10728.

Pancer, Z., Amemiya, C. T., Ehrhardt, G. R., Ceitlin, J., Gartland, G. L., & Cooper, M. D. (2004). Somatic diversification of variable lymphocyte receptors in the agnathan sea lamprey. Nature, 430, 174–180.

Patel, B., Banerjee, R., Basu, M., Lenka, S., Samanta, M., & Das, S. (2016). Molecular cloning of IgZ heavy chain isotype in Catla catla and comparative expression profile of IgZ and IgM following pathogenic infection. Microbiology and immunology, 60, 561–567.

Peterson, P. A., Cunningham, B. A., Berggård, I., & Edelman, G. M. (1972). *β*2-Microglobulin—a free immunoglobulin domain. Proceedings of the National Academy of Sciences, 69, 1697–1701.

Piazzon, M. C., Galindo-Villegas, J., Pereiro, P., Estensoro, I., Calduch-Giner, J. A., Gómez-Casado, E., Novoa, B., Mulero, V., Sitja`-Bobadilla, A., & Pérez-Sánchez, J. (2016). Differential modulation of IgT and IgM upon parasitic, bacterial, viral, and dietary challenges in a Perciform fish. Frontiers in immunology, 7, 637.

Price, M. N., Dehal, P. S., & Arkin, A. P. (2010). Fasttree 2–approximately maximum-likelihood trees for large alignments. PloS one, 5.

Ross, D. A., Wilson, M. R., Miller, N. W., Clem, L. W., Warr, G. W., Ross, D. A., Wilson, M. R., Miller, N. W., Clem, L. W., & Warr, G. W. (1998). Evolutionary variation of immunoglobulin *μ* heavy chain RNA processing pathways: origins, effects, and implications. Immunological reviews, 166, 143–151.

Ryo, S., Wijdeven, R. H., Tyagi, A., Hermsen, T., Kono, T., Karunasagar, I., Rombout, J. H., Sakai, M., Verburg-van Kemenade, B. L., & Savan, R. (2010). Common carp have two subclasses of bonyfish specific antibody IgZ showing differential expression in response to infection. Developmental & Comparative Immunology, 34, 1183–1190.

Sahoo, M., Edholm, E.-S., Stafford, J. L., Bengtén, E., Miller, N. W., & Wilson, M. (2008). B cell receptor accessory molecules in the channel catfish, Ictalurus punctatus. Developmental & Comparative Immunology, 32, 1385–1397.

Savan, R., Aman, A., Nakao, M., Watanuki, H., & Sakai, M. (2005a). Discovery of a novel immunoglobulin heavy chain gene chimera from common carp (Cyprinus carpio L.). Immunogenetics, 57, 458–463.

Savan, R., Aman, A., Sato, K., Yamaguchi, R., & Sakai, M. (2005b). Discovery of a new class of immunoglobulin heavy chain from fugu. European journal of immunology, 35, 3320–3331.

Sievers, F., & Higgins, D. G. (2014). Clustal Omega, accurate alignment of very large numbers of sequences. In Multiple sequence alignment methods (pp. 105–116). Springer.

Star, B., Nederbragt, A. J., Jentoft, S., Grimholt, U., Malmstrøm, M., Gregers, T. F., Rounge, T. B., Paulsen, J., Solbakken, M. H., Sharma, A. et al. (2011). The genome sequence of Atlantic cod reveals a unique immune system. Nature, 477, 207.

Stenvik, J., & Jørgensen, T. Ø. (2000). Immunoglobulind (igd) of atlantic cod has a unique structure. Immunogenetics, 51, 452–461.

Sun, S.-C., Lindstrom, I., Boman, H. G., Faye, I., & Schmidt, O. (1990). Hemolin: an insect-immune protein belonging to the immunoglobulin superfamily. Science, 250, 1729–1732.

Sun, Y., Huang, Y., Li, X., Baldwin, C. C., Zhou, Z., Yan, Z., Crandall, K. A., Zhang, Y., Zhao, X., Wang, M. et al. (2016). Fish-t1k (Transcriptomes of 1,000 Fishes) Project: large-scale transcriptome data for fish evolution studies. Gigascience, 5, 18.

Tang, X., Du, Y., Sheng, X., Xing, J., & Zhan, W. (2018). Molecular cloning and expression analyses of immunoglobulin tau heavy chain (IgT) in turbot, Scophthalmus maximus. Veterinary immunology and immunopathology, 203, 1–12.

Tian, J., Sun, B., Luo, Y., Zhang, Y., & Nie, P. (2009). Distribution of IgM, IgD and IgZ in mandarin fish, Siniperca chuatsi lymphoid tissues and their transcriptional changes after Flavobacterium columnare stimulation. Aquaculture, 288, 14–21.

Velázquez, J., Acosta, J., Lugo, J. M., Reyes, E., Herrera, F., González, O., Morales, A., Carpio, Y., & Estrada, M. P. (2018). Discovery of immunoglobulin T in nile tilapia (Oreochromis niloticus): A potential molecular marker to understand mucosal immunity in this species. Developmental & Comparative Immunology, 88, 124–136.

Waterhouse, A. M., Procter, J. B., Martin, D. M., Clamp, M., & Barton, G. J. (2009). Jalview version 2—a multiple sequence alignment editor and analysis workbench. Bioinformatics, 25, 1189–1191.

Wilson, M., Bengtén, E., Miller, N. W., Clem, L. W., Du Pasquier, L., & Warr, G. W. (1997). A novel chimeric ig heavy chain from a teleost fish shares similarities to IgD. Proceedings of the National Academy of Sciences, 94, 4593–4597.

Xia, H., Liu, W., Wu, K., Wang, W., & Zhang, X. (2016). sIgZ exhibited maternal transmission in embryonic development and played a prominent role in mucosal immune response of Megalabrama amblycephala. Fish & shellfish immunology, 54, 107–117.

Xiao, F., Wang, Y., Yan, W., Chang, M., Yao, W., Xu, Q., Wang, X., Gao, Q., & Nie, P. (2010). Ig heavy chain genes and their locus in grass carp Ctenopharyngodon idella. Fish & shellfish immunology, 29, 594–599.

Xu, J., Yu, Y., Huang, Z., Dong, S., Luo, Y., Yu, W., Yin, Y., Li, H., Liu, Y., Zhou, X. et al. (2019). Immunoglobulin (Ig) heavy chain gene locus and immune responses upon parasitic, bacterial and fungal infection in loach, Misgurnus anguillicaudatus. Fish & shellfish immunology, 86, 1139–1150.

Yasuike, M., De Boer, J., von Schalburg, K. R., Cooper, G. A., McKinnel, L., Messmer, A., So, S., Davidson, W. S., & Koop, B. F. (2010). Evolution of duplicated IgH loci in atlantic salmon, salmo salar. BMC genomics, 11, 486.

Ye, J., Bromage, E. S., & Kaattari, S. L. (2010). The strength of B cell interaction with antigen determines the degree of IgM polymerization. The journal of immunology, 184, 844–850.

Zhang, N., Zhang, X.-J., Chen, D.-D., Sunyer, J. O., & Zhang, Y.-A. (2017). Molecular characterization and expression analysis of three subclasses of IgT in rainbow trout (Oncorhynchus mykiss). Developmental & Comparative Immunology, 70, 94–105.

Zhang, S.-M., Adema, C. M., Kepler, T. B., & Loker, E. S. (2004). Diversification of Ig superfamily genes in an invertebrate. Science, 305, 251–254.

Zhang, Y.-A., Salinas, I., & Sunyer, J. O. (2011). Recent findings on the structure and function of teleost IgT. Fish & shellfish immunology, 31, 627–634.

Zhao, Y., Cui, H., Whittington, C. M., Wei, Z., Zhang, X., Zhang, Z., Yu, L., Ren, L., Hu, X., Zhang, Y. et al. (2009). Ornithorhynchus anatinus (platypus) links the evolution of immunoglobulin genes in eutherian mammals and nonmammalian tetrapods. The Journal of Immunology, 183, 3285–3293.

